# Reovirus σ3 protein limits interferon expression and cell death induction

**DOI:** 10.1101/2020.07.19.210872

**Authors:** Katherine E Roebke, Yingying Guo, John S. L. Parker, Pranav Danthi

## Abstract

Induction of necroptosis by mammalian reovirus requires both type I interferon (IFN)-signaling and viral replication events that lead to production of progeny genomic dsRNA. The reovirus outer capsid protein µ1 negatively regulates reovirus-induced necroptosis by limiting RNA synthesis. To determine if the outer capsid protein σ3, which interacts with µ1, also functions in regulating cell death, we used siRNA-mediated knockdown. Similar to that observed by diminishment of µ1 expression, knockdown of newly synthesized σ3 enhances necroptosis. σ3 knockdown does not impact reovirus RNA synthesis. Instead, this increase in necroptosis following σ3 knockdown is accompanied by an increase in IFN production. Furthermore, ectopic expression of σ3 is sufficient to block IFN expression following infection. Surprisingly, the capacity of σ3 protein to bind dsRNA does not impact its capacity to diminish production of IFN. Consistent with this, infection with a virus harboring a mutation in the dsRNA binding domain of σ3 does not result in enhanced production of IFN or cell death. Together, these data suggest that σ3 limits the production of IFN to control innate immune signaling and cell death following infection through a mechanism that is independent of its dsRNA binding capacity.

**IMPORTANCE:** We use mammalian reovirus as a model to study how virus infection modulates innate immune signaling and cell death induction. Here we sought to determine how viral factors regulate these processes. Our work highlights a previously unknown role for the reovirus outer capsid protein σ3 in limiting the induction of a necrotic form of cell death called necroptosis. Induction of cell death by necroptosis requires production of interferon. σ3 limits the induction of necroptosis by preventing excessive production of interferon following infection.

## INTRODUCTION

Innate immune signaling following virus infection is a common mechanism by which cells can limit viral proliferation and spread. A key component of this innate immune signaling is interferon (IFN) (1). Virus-induced activation of interferon (IFN) signaling leads to the expression of hundreds of target genes called interferon stimulated genes (ISGs) that help establish an antiviral state and curb virus replication (2). ISGs can also participate in the induction of various forms of cell death by leading to the expression of death ligands and cytokines or by altering key host processes (2, 3). To ensure productive infection, viruses have evolved strategies to prevent IFN expression, block IFN signaling, or interfere with the function of specific ISGs that control cell death (4).

Mammalian orthoreovirus (reovirus) is used as a model to investigate virus induced cell death. Reovirus is a dsRNA virus that replicates in the cytoplasm (5). Reovirus induces two forms of cell death, apoptosis and necroptosis (6, 7). The execution of these two forms of death require distinct sets of proteins. Among these, type I IFN expression and signaling is required for reovirus-induced necroptosis but not apoptosis (8, 9). To induce IFN-dependent necroptosis, the incoming reovirus dsRNA genome is detected by cellular sensors retinoic acid-inducible gene I (RIG-I) and melanoma differentiation-associated protein 5 (MDA5)(9). These sensors signal via the mitochondrial antiviral-signaling protein (MAVS) to promote the expression of type I interferon. It is deduced that signaling via MAVS-dependent IFN expression requires the phosphorylation and nuclear translocation of the IRF3 transcription factor. IFN is released from infected cells and signals via the IFN α/β receptor (IFNAR) to induce cell death (9). Together with events dependent on synthesis of viral progeny dsRNA, IFN signaling leads to activation of RIP1 and RIP3-dependent necroptosis which is likely mediated by the mixed lineage kinase domain-like (MLKL) effector protein. It is expected that viral factors that influence levels of IFN produced, those that affect signaling by IFN, and those that modulate the activity of the death inducing ISGs will likely impact the induction of cell death by reovirus. However, reovirus proteins with such functions have not been identified.

We recently identified a role for the reovirus µ1 protein in regulating viral replication and cell death (10). Knockdown of this essential outer capsid protein with small interfering RNA (siRNA) results in enhanced viral gene expression, protein production, and minus strand RNA synthesis to generate new genome. These phenotypes are accompanied by an increase in necroptosis. During reovirus replication, viral gene transcription and minus strand RNA synthesis occur within core particles (5). Because dsRNA itself or a product dependent on dsRNA synthesis is required for induction of necroptosis (9), we propose that µ1, by virtue of its capacity to assemble on transcribing core particles in the cytoplasm of host cells, limits cell death by shutting off viral transcription. µ1 associates with another viral outer capsid protein σ3 to form heterohexamers prior to assembly on the core particles (11-13). Thus, it might be expected that knockdown of σ3 also may produce a similar effect on accumulation of viral RNAs and cell death. Data supporting this idea are thus far lacking. Evaluating how σ3 affects cell death was a goal of our study here.

In addition to coassembling with µ1, σ3 has other functions in the cell. σ3 affects capsid stability and cell entry kinetics when assembled on virus particles (14-17). A function is also attributed to newly synthesized, intracellular σ3. σ3 has a dsRNA binding domain, which interacts with dsRNA in a sequence-independent manner, and prevents the activation of the dsRNA-activated protein kinase R (PKR) (18-20). PKR is an ISG which phosphorylates the eukaryotic translation initiation factor 2A (eIF2α), thereby causing translational arrest (21). σ3 competes for dsRNA binding with PKR and suppresses the effect on translation to promote viral protein synthesis and productive infection (22, 23). The dsRNA binding capacity of σ3 is abrogated by its interaction with outer capsid protein µ1 (11, 23). These additional functions of σ3 could also therefore have an effect on cell death.

In this study, we aimed to determine if σ3 regulates cell death. We assessed this using siRNA-mediated knockdown of newly synthesized σ3. Knockdown of σ3 increases necroptosis in L929 cells, while not impacting apoptotic cell death in HeLa cells. Knockdown of σ3 does not alter levels of viral mRNA or dsRNA genome. Instead, we observe an increase in IFN-β production and signaling. The effect of σ3 on IFN-β expression is not related to its capacity to bind dsRNA. These studies uncover a new function for the σ3 protein and indicate how σ3 levels can modulate the host innate immune response and cell death.

## MATERIALS AND METHODS

### Cells

L929 cells obtained from ATCC were maintained in Eagle’s MEM (Lonza) supplemented to contain 5% fetal bovine serum (FBS) (Life Technologies), and 2 mM L-glutamine (Invitrogen). Spinner-adapted Murine L929 cells were maintained in Joklik’s MEM (Lonza) supplemented to contain 5% fetal bovine serum (FBS) (Sigma-Aldrich), 2 mM L-glutamine (Invitrogen), 100 U/ml penicillin (Invitrogen), 100 μg/ml streptomycin (Invitrogen), and 25 ng/ml amphotericin B (Sigma-Aldrich). Spinner-adapted L929 cells were used for cultivating, purifying and titering viruses.

HeLa cells and HEK293 cells obtained from Melanie Marketon’s lab at Indiana University were maintained in Dulbecco’s MEM (Lonza) supplemented to 5% FBS (Sigma-Aldrich, and 2mM L-glutamine (Invitrogen).

### Reagents

Q-VD-OPh was purchased from Cayman Chemical Company. Pyridone 6 was purchased from Fisher Scientific. Custom synthesized siRNAs were purchased from Dharmacon. siRNA targeting β-galactosidase was used as control. The siRNA sequences used are as follows: β-galactosidase – CUACACAAAUCAGCGAUUU, σ3 – GCGCAAGAGGGAUGGGACA, and RIP3 – GGUAAAGCAUUAUCUGUCU. Antisera raised against reovirus capsid was obtained from T. Dermody and have been previously described (24). Monoclonal 4F2 antibody against σ3 was obtained from Iowa hybridoma bank (25). Monoclonal antibody against IFNAR and rabbit antisera against RIP3 were purchased from Santa Cruz Biotechnology and Pro-science respectively. Mouse antiserum specific for PSTAIR was purchased from Sigma. Alexa Fluor-conjugated or IR dye conjugated anti-mouse IgG and anti-rabbit IgG secondary antibodies were purchased from Invitrogen or Licor.

### Virus purification

Prototype reovirus strains of T3D, T1L, and σ3 mutants were regenerated by plasmid based reverse genetics (26). Purified reovirus virions were generated using second- or third-passage L-cell lysates stocks of reovirus. Viral particles were Vertrel-XF (Dupont) extracted from infected cell lysates, layered onto 1.2- to 1.4-g/cm^3^ CsCl gradients, and centrifuged at 187,183 x *g* for 4 h. Bands corresponding to virions (1.36 g/cm^3^) were collected and dialyzed in virion-storage buffer (150 mM NaCl, 15 mM MgCl_2_, 10 mM Tris-HCl [pH 7.4]) (27). UV-inactivated virus was generated by using a UV cross-linker (CL-1000 UV cross-linker; UVP). Virus diluted in phosphate-buffered saline (PBS) was placed into a 60-mm tissue culture dish and irradiated with short-wave (254-nm) UV at a distance of 10 cm for 1 min at 120,000 µJ/cm^2^.

### Plaque assays

Plaque assays to determine infectivity were performed as previously described with some modifications (27). Briefly, control or heat-treated virus samples were diluted into PBS supplemented with 2 mM MgCl_2_ (PBS_Mg_). L cell monolayers in 6-well plates (Greiner Bio-One) were infected with 100 μl of diluted virus for 1 h at room temperature. Following the viral attachment incubation, the monolayers were overlaid with 4 ml of serum-free medium 199 (Sigma-Aldrich) supplemented with 1% Bacto Agar (BD Biosciences), 10 μg/ml TLCK-treated chymotrypsin (Worthington, Biochemical), 2 mM L-glutamine (Invitrogen), 100 U/ml penicillin (Invitrogen), 100 μg/ml streptomycin (Invitrogen), and 25 ng/ml amphotericin B (Sigma-Aldrich). The infected cells were incubated at 37°C, and plaques were counted 5 d post infection.

### siRNA transfection

In 96-well plates, 0.25 μl Lipofectamine 2000 was used to transfect siRNA to a final concentration of 100 nM, or 0.75µl INTERFERin (Polyplus) was used to transfect siRNA to a final concentration of 5 nM. Cells (1 × 10^4^) were seeded on top of the siRNA-lipofectamine/INTERFERin mixture. In 24-well plates, 1.5 µl Lipofectamine 2000 was used to transfect siRNA to a final concentration of 42 nM or 3 µl of INTERFERin was used to transfect siRNA to a final concentration of 15 nM. Cells (1 × 10^5^) were seeded on top of the siRNA-lipofectamine/INTERFERin mixture. Virus infection was performed 28-48 h following siRNA transfection.

### Plasmid construction and transfections

pCMV-S4 encoding σ3 was obtained from the T. Dermody laboratory (24). Site-directed mutagenesis of the pCMV-S4 vector was performed using protocols and materials from QuickChange Lightning mutagenesis kit (Agilent). Nearly confluent monolayers of HEK293 cells in 12 well plates were transfected with either 0.5 μg of empty vector or 0.25 μg each of WT σ3 or σ3 K293T expression vector using 1.5 µl Lipofectamine 2000 according to the manufacturer’s instructions. Transfected cells were incubated at 37°C for 24 h prior to infection to allow expression from the plasmids.

### Infection and preparation of extracts

Cells were either adsorbed with PBS or T3D at room temperature for 1 h, followed by incubation with media at 37°C for the indicated time interval. When needed, DMSO, Q-VD-OPh, anti-IFNAR Ab, or Pyridone6 were added to the media immediately after the 1 h adsorption period. For preparation of whole cell lysates, cells were washed in phosphate-buffered saline (PBS) and lysed with 1X RIPA (50 mM Tris [pH 7.5], 50 mM NaCl, 1% TX-100, 1% DOC, 0.1% SDS, and 1 mM EDTA) containing a protease inhibitor cocktail (Roche) and 2 mM PMSF, followed by centrifugation at 15000 × *g* for 10 min to remove debris.

### Immunoblotting

Cell lysates were resolved by electrophoresis on 10% polyacrylamide gels and transferred to nitrocellulose membranes. Membranes were blocked for at least 1 h in TBS containing T20 Starting Block (Thermo Fisher) and incubated with antisera against reovirus (1:5000), σ3 (1:1000), RIP3 (1:1000), or PSTAIR (1:10000) at 4°C overnight. Membranes were washed three times for 5 min each with washing buffer (TBS containing 0.1% Tween-20) and incubated with 1:20000 dilution of Alexa Fluor conjugated goat anti-rabbit IgG (for reovirus polyclonal and RIP3) or goat anti-mouse IgG (for PSTAIR) in blocking buffer. Following three washes, membranes were scanned using an Odyssey Infrared Imager (LI-COR) or Chemidoc MP Imaging System (Biorad). Band intensity was analyzed using Image Studio Lite software (LICOR).

### Quantitation of cell death by Acridine orange-ethidium bromide (AOEB) staining

ATCC L929 cells or HeLa cells (2×10^5^) grown in 96 well-plates were siRNA transfected as described above and adsorbed with the indicated MOI of reovirus at room temperature for 1 h. The percentage of dead cells at the indicated time following infection was determined using AOEB staining as described previously (6). For each experiment, >200 cells were counted, and the percentage of isolated cells exhibiting orange staining (EB positivity) was determined by epi-illumination fluorescence microscopy using a fluorescein filter set on an Olympus IX71 microscope.

### Assessment of caspase-3/7 activity

ATCC L929 cells (1 × 10^4^) were siRNA transfected as described above and seeded into black clear-bottom 96-well plates. 24 h following transfection, cells were adsorbed with 10 PFU/cell of reovirus in serum-free medium at room temperature for 1 h. Following incubation of cells at 37°C for 48 h, caspase-3/7 activity was quantified using the Caspase-Glo-3/7 assay system (Promega). Luminescence was quantified using plate reader Synergy H1 Hybrid Reader (BioTek).

### RT-qPCR

RNA was extracted from infected cells at various time intervals after infection using Total RNA mini kit (Biorad). For RT-qPCR, 0.5 to 2 µg of RNA was reverse transcribed with the High Capacity cDNA Reverse Transcription Kit (Applied Biosystems) using random hexamers for amplification of cellular genes or gene specific primers for amplification of either minus or plus strand viral RNA. Gene specific primers were used at a final concentration of 0.1 µM. A 1:10 dilution of the cDNA was subjected to PCR using SYBR Select Master Mix (Applied Biosystems). Fold increase in gene expression with respect to control sample (indicated in each figure legend) was measured using the ΔΔC_T_ method (28). Calculations for determining ΔΔC_T_ values and relative levels of gene expression were performed as follows: (i) fold increase in cellular gene expression = 2 _-[(gene of interest CT – GAPDH CT)T3D infected – (gene of interest CT – GAPDH CT)Mock infected]_; (ii) fold increase in viral gene expression = 2 _-[(T3D S1 CT – GAPDH CT)σ3 siRNA – (T3D S1 CT – GAPDH CT)control siRNA]_

### Assessment of IFNβ levels by ELISA

ATCC L929 cells (2 × 10^5^) grown in 24 well-plates were siRNA transfected as described above and adsorbed with the indicated MOI of reovirus at room temperature for 1 h. Media was harvested at indicated times post infection and IFNβ levels were quantified using a standard curve via Mouse IFN-beta DuoSet ELISA kit (R&D systems) using manufacturer’s instructions. Optical density was quantified using plate reader Synergy H1 Hybrid Reader (BioTek).

### Statistical analysis

Statistical significance between experimental groups was determined using the unpaired student’s *t*-test function in excel and graphed using Graphpad Prism software. Statistical analyses for differences in gene expression by RT-qPCR were done on the ΔC_T_ values.

## RESULTS

### Knockdown of newly synthesized σ3 drives an increase in cell death in L929 cells

The levels of outer capsid protein µ1 modulate cell death (10). Our work indicates that newly synthesized µ1 negatively regulates the induction of necroptosis in L929 cells by preventing excessive accumulation of viral gene products. In this study, we aim to determine if outer capsid protein σ3, which interacts with µ1, also regulates cell death. To determine if σ3 modulates induction of cell death, we knocked down the levels of the σ3 protein in L929 cells infected with reovirus type 3 prototype strain T3D using siRNA. This strain has been extensively characterized for its capacity to induce apoptosis or necroptosis, depending on the cell type (6, 9). As expected, the siRNA significantly diminished the levels of newly synthesized σ3 in infected cells (Figure 1A). To assess the impact of σ3 knockdown on levels of cell death, L929 cells transfected with either control or σ3 siRNA were infected with T3D for 24 h and cell death was quantified by acridine orange-ethidium bromide (AOEB) staining (Figure 1B). Acridine orange, a membrane-permeable dye, stains all cells. In contrast, ethidium bromide incorporates into and stains only the DNA of cells with compromised membranes, such as following cell death. Diminishment of σ3 expression resulted in increased cell death in infected L929 cells.

**Figure 1.**
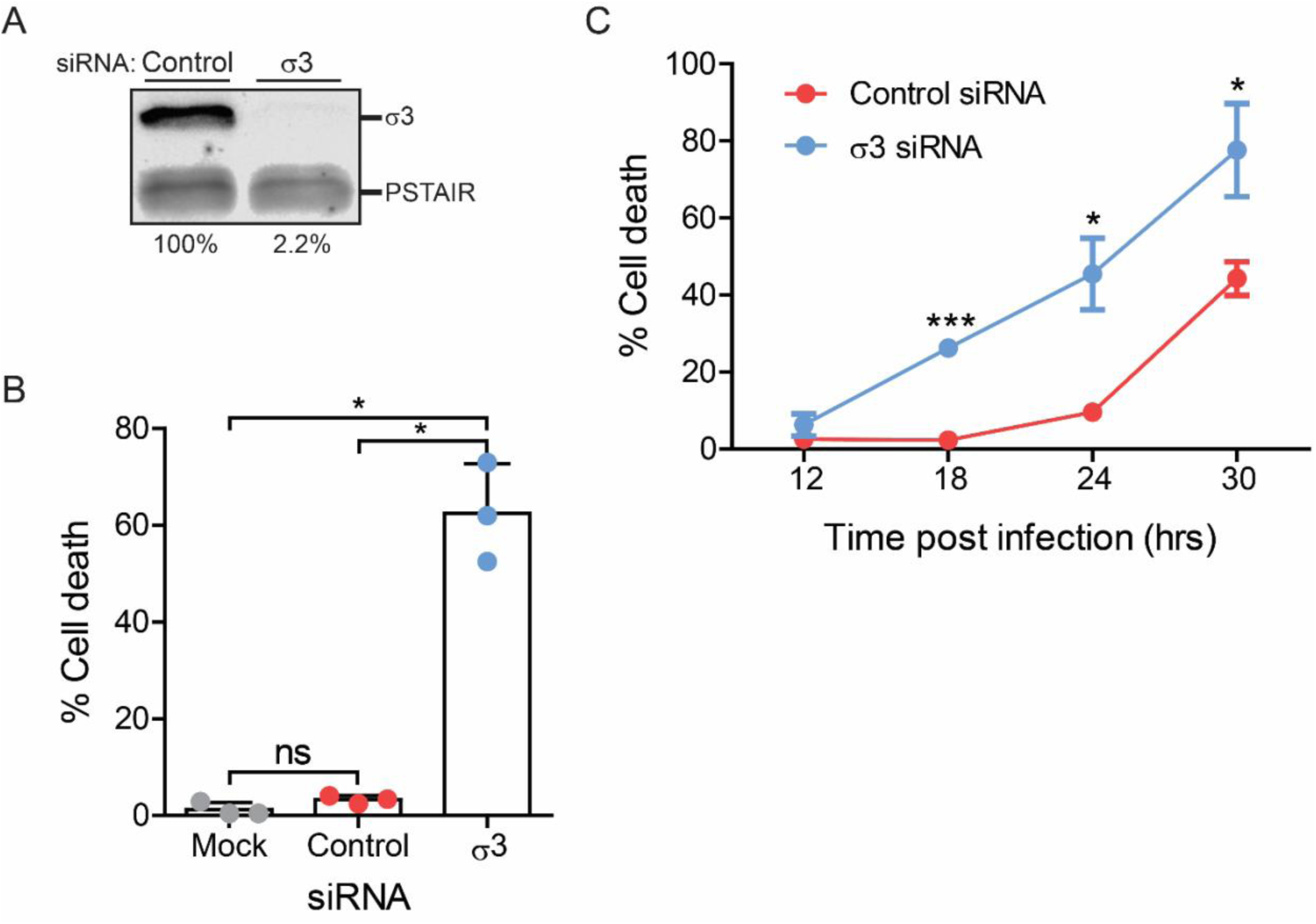
Knockdown of newly synthesized σ3 increases cell death in L929 cells. L929 cells were transfected with either control β-gal or σ3 siRNA using Lipofectamine 2000. 24 h following transfection, the cells were infected with T3D at an MOI of 10 PFU/cell and processed as described. (A) Cell lysates prepared 24 h following infection were immunoblotted for the σ3 protein using anti-reovirus antisera and anti-PSTAIR mAb. The level of σ3 relative to PSTAIR in control siRNA treated cells was considered 100%. (B) Cell death was quantified 24 h following infection by AOEB staining. (C) Cell death was quantified at the indicated times by AOEB staining. (B and C) Cell death for three independent infections and mean are shown. Error bars indicate SD. P values were determined by student’s t-test. *, P<0.05.

Relative to control siRNA-treated cells which exhibited only minimal levels of cell death (∼3%) at 24 h following reovirus infection, knockdown of σ3 resulted in significantly more cell death (∼60%). When cell death was quantified over time, we observed that cells transfected with control siRNA started exhibiting cell death around 24-30 h post infection (Figure 1C). In contrast, cells transfected with σ3 siRNA succumbed to reovirus infection as early as 18 h post infection. These data suggest that newly synthesized σ3 negatively regulates cell death in L929 cells following reovirus infection. L929 cells infected with reovirus undergo RIP3-dependent necroptosis (9, 10). It is possible, however, that σ3 knockdown changes the nature of cell death in these cells. To determine what form of cell death is enhanced by knockdown of σ3 following T3D infection, we quantified cell death in σ3 knockdown cells treated with inhibitors that block either apoptotic or necroptotic cell death. Although treatment of cells with the pan-caspase inhibitor, QVD, completely blocked effector caspase (caspase-3/7) activity in reovirus-infected cells (Figure 2A), it did not influence cell death in the presence of σ3 siRNA knockdown (Figure 2B). These data suggest that, following knockdown of σ3, reovirus continues to induce a nonapoptotic form of cell death. To block necroptosis, we used a siRNA specific to the necroptotic kinase, RIP3 (Figure 2C). Simultaneous knockdown of both σ3 and RIP3 resulted in a diminishment of cell death (Figure 2D). Thus, even following knockdown of σ3, L929 cells infected with reovirus undergo cell death via necroptosis.

**Figure 2.**
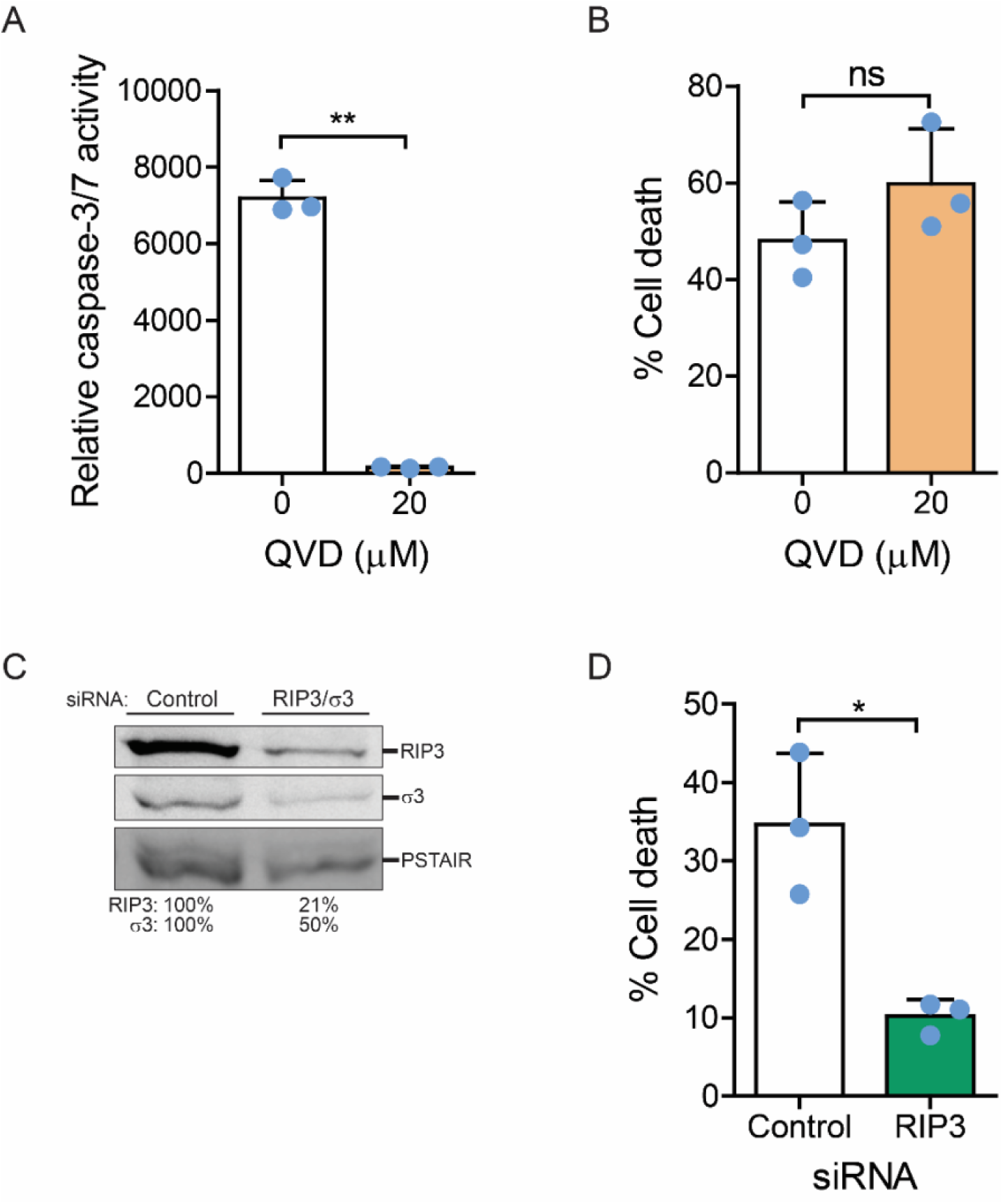
σ3 knockdown increases necroptotic cell death in L929 cells. L929 cells were transfected with σ3 siRNA using (A, C-D) INTERFERin or (B) Lipofectamine 2000. 24 h following transfection, the cells were infected with T3D at an MOI of 10 PFU/cell and either left untreated or were treated with 20 µM Q-VD-OPh when infection was initiated. At 30 h following infection, (A) caspase-3/7 activity was determined by chemiluminescent assay or (B) cell death was quantified by AOEB staining. (C and D) L929 cells were cotransfected with siRNA specific to σ3 and β-gal or σ3 and RIP3 using INTERFERin. (C) Cell lysates prepared 24 h following transfection were immunoblotted using anti-reovirus, anti-RIP3, and anti-PSTAIR antibodies. The level of σ3 and RIP3 relative to PSTAIR in control siRNA treated cells was considered 100%. (D) 24 h following transfection cells were infected with T3D at an MOI of 10 PFU/cell. At 30 h following infection cell death was quantified by AOEB staining. Mean values for three independent infections are shown. Error bars indicate SD. P values were determined by student’s t-test. *, P<0.05; **, P<0.005.

### Newly synthesized σ3 does not function in reovirus-induced apoptotic cell death

HeLa cells do not express the RIP3 protein (29), and thus primarily undergo apoptotic cell death following reovirus infection (10). To determine whether σ3 impacts apoptosis induced by reovirus, we knocked down the levels of σ3 in reovirus-infected HeLa cells using siRNA. Analogous to what we observed in L929 cells, the siRNA significantly diminished levels of newly synthesized σ3 in T3D infected HeLa cells (Figure 3A). To determine whether knockdown of σ3 affects the levels of cell death, HeLa cells transfected with control or σ3 siRNA were infected with T3D and cell death was quantified by AOEB staining (Figure 3B). In contrast to what we observed in L929 cells, diminishment in σ3 expression in HeLa cells did not enhance cell death. These data indicate that newly synthesized σ3 does not affect the efficiency of apoptotic cell death following reovirus infection.

**Figure 3.**
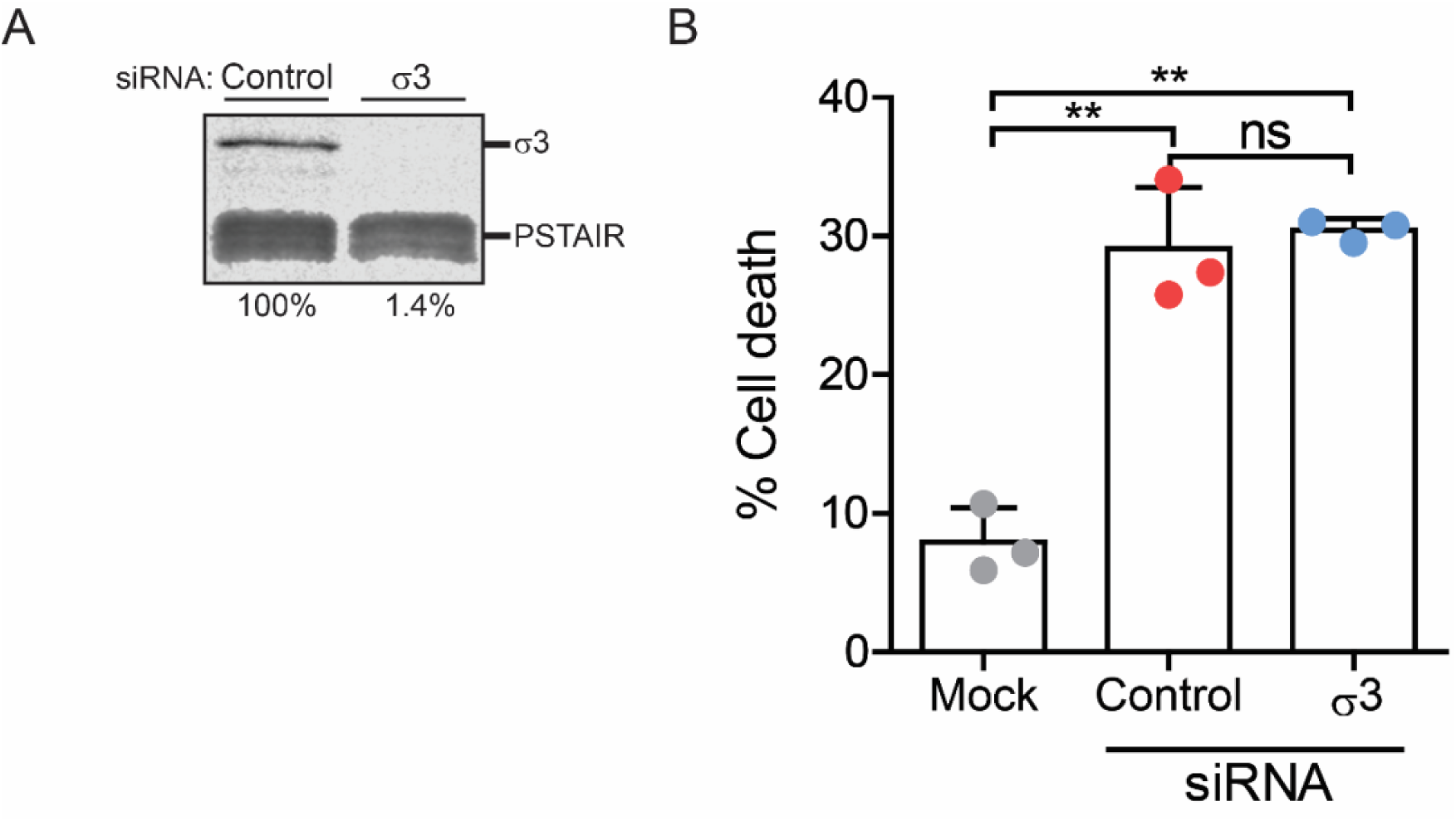
Newly synthesized σ3 does not impact apoptotic cell death. HeLa cells were transfected with either control β-gal or σ3 siRNA using INTERFERin. 24 h following transfection, the cells were infected and processed as described. (A) Cells were infected with T3D at an MOI of 10 PFU/cell. Cell lysates prepared 24 h following infection were immunoblotted for the σ3 protein using anti-reovirus antisera and anti-PSTAIR mAb. Levels of σ3 bands relative to PSTAIR are indicated. The level of σ3 relative to PSTAIR in control siRNA treated cells was considered 100%. (B) Cells were either mock infected or infected with T3D at an MOI of 50 PFU/cell. At 44 h post infection, cell death was quantified by AOEB staining. Mean values for three independent infections are shown. Error bars indicate SD. P values were determined by student’s t-test. ** p < 0.005

### Loss of σ3 does not impact replication events

Our recent work studying the role of µ1 in replication and cell death demonstrated that this outer capsid protein limits the cell death response by negatively regulating viral replication events (10). We observed that knockdown of newly synthesized µ1 results in increased production of secondary transcripts and dsRNA genome. To determine if the increase in cell death observed following knockdown of σ3 is also due to enhanced production of viral dsRNA genome and mRNA, we quantified levels of viral minus-strand and plus-strand RNA respectively in infected cells by strand-specific RT and qPCR (Figure 4). We used S1 gene-derived RNAs as representatives for these experiments. In contrast to what was observed following knockdown of µ1, loss of σ3 does not drive any significant change in minus or plus strand RNA relative to control siRNA-treated cells following infection. Knockdown of µ1 also leads to an increased accumulation of other viral proteins.

**Figure 4.**
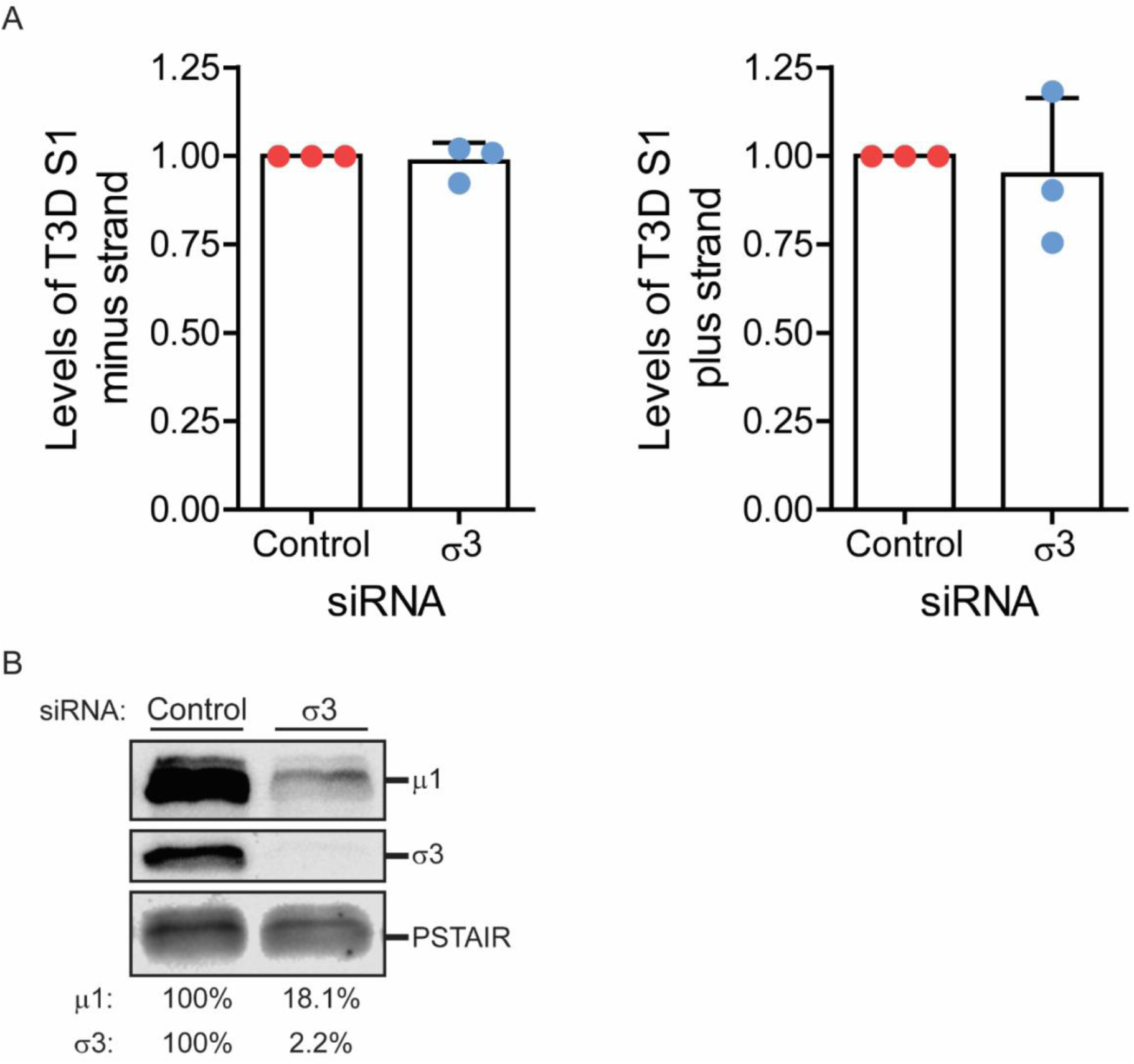
Levels of viral plus and minus strand RNA are not altered following knockdown of σ3. L929 cells were transfected with control β-gal or σ3 siRNA using INTERFERin. 24 h following transfection, cells were infected with T3D at an MOI of 10 PFU/cell. (A) RNA extracted from infected cells harvested at 21 h post infection was reverse transcribed using primers specific to the minus strand or plus strand of the T3D S1 gene or GAPDH mRNA. Levels of accumulated T3D S1 (left) minus strand or (right) plus strand RNA relative to GAPDH mRNA was quantified by qPCR and comparative C_T_ analysis. The ratio of T3D S1 RNA to GAPDH in control siRNA-treated cells was set to 1. Mean values for three independent infections are shown. Error bars indicate SD. (B) Cell lysates prepared 24 h following infection were immunoblotted for the µ1 and σ3 protein using anti-reovirus antisera and anti-PSTAIR mAb. Levels of µ1 and σ3 bands relative to PSTAIR are indicated. The level of µ1 and σ3 relative to PSTAIR in control siRNA treated cells was considered 100%.

While we think the increase in viral protein expression following diminishment of µ1 expression is a result of greater steady state levels of viral mRNAs, we tested the effect of σ3 knockdown on the levels of other viral proteins. We found that upon knockdown of σ3, the expression of µ1 and other viral proteins (Figure 4B and data not shown) was also decreased. These data suggest that the mechanism driving increased cell death following σ3 knockdown is not through an increase in accumulation of viral gene products, and is thus different than that of µ1 knockdown.

### Reovirus-induced necroptosis is less sensitive to blockade of IFN signaling following knockdown of σ3

Previous work from our laboratory demonstrated that necroptosis following reovirus infection requires type I IFN expression and signaling via the IFN-α/β receptor (IFNAR)(9, 10). Thus, treatment with a monoclonal antibody against IFNAR (30) diminishes cell death in T3D infected L929 cells. To determine if necroptosis following σ3 knockdown is also affected by IFNAR, we quantified cell death following reovirus infection in L929 cells treated with anti-IFNAR Ab (Figure 5). As expected, in control-siRNA treated cells, treatment with the anti-IFNAR Ab significantly reduced cell death by ∼75%. In contrast, following loss of σ3 expression the anti-IFNAR antibody did not reduce cell death. As an alternative method to block IFN signaling, we used pyridone-6, which targets the downstream, JAK kinase (31). As expected, control-siRNA treated cells infected with reovirus in the presence of the JAK inhibitor pyridone-6 exhibited ∼50% less cell death than untreated cells (Figure 5). Analogous to what was observed following treatment with the anti-IFNAR antibody, treatment with the JAK inhibitor did not reduce cell death following σ3 knockdown. Higher concentration of IFNAR Ab or the JAK inhibitor were either not possible to use or compromised cell health. Together, these data suggest that loss of σ3 expression results in decreased sensitivity of reovirus-induced necroptosis to blockade of IFN signaling.

**Figure 5.**
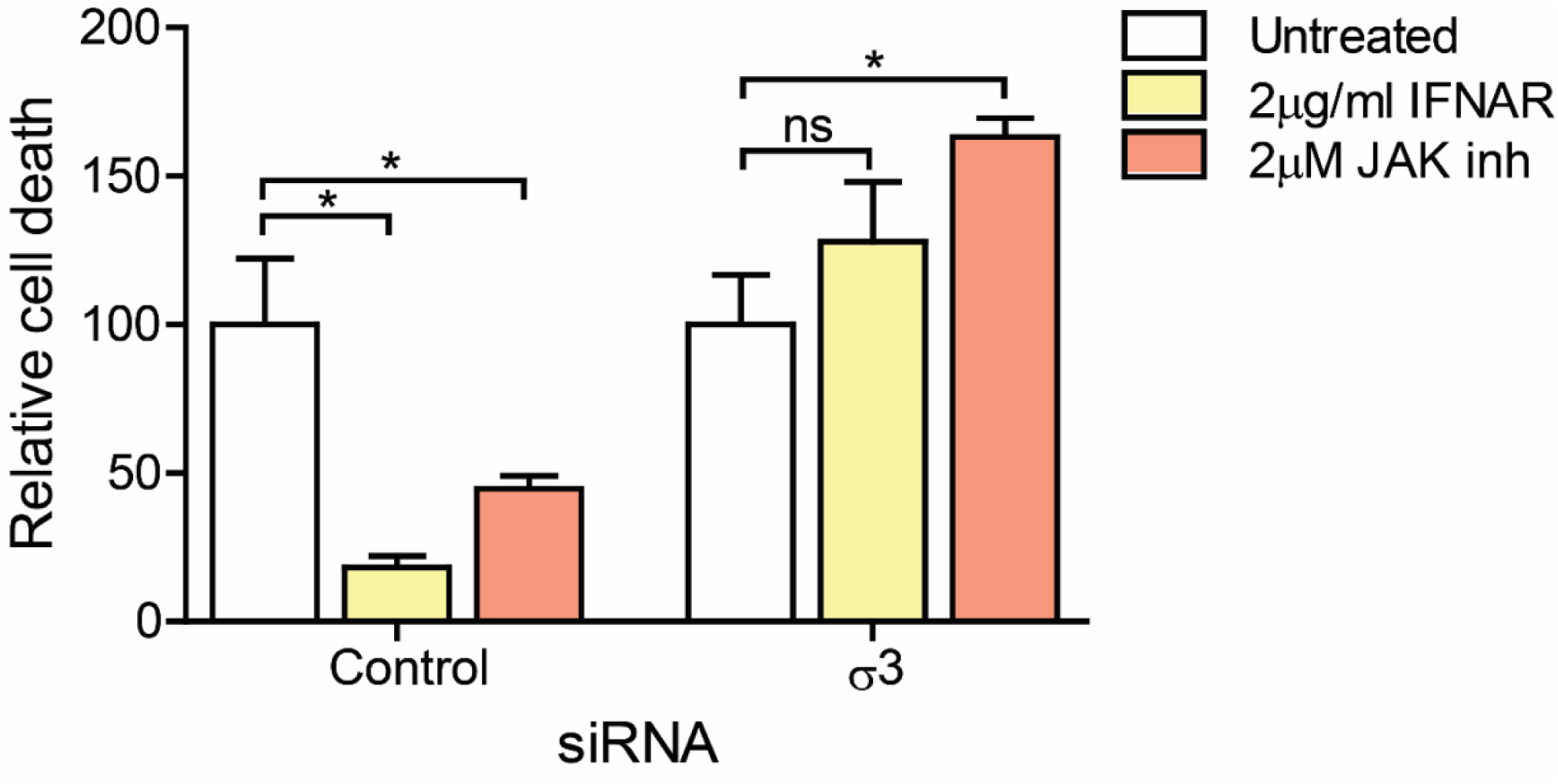
Cell death induced following σ3 knockdown is not sensitive to blockade of type I IFN signaling. L929 cells were transfected with control β-gal or σ3 siRNA using INTERFERin. 24 h following transfection the cells were infected with T3D at an MOI of 10 PFU/cell and either left untreated or were treated with 2µg/ml anti-IFNAR mAb or 2µM Pyridone 6 when infection was initiated. At 42 h following infection, cell death was quantified by AOEB staining. Percent decrease in cell death in the presence of treatment was quantified for three independent infections. Cell death in the absence of treatment was set to 100 for each siRNA-treated sample. Error bars indicate SD. P values were determined by student’s t-test. *, P<0.05.

### IFN signaling is increased when σ3 is knocked down

The anti-IFNAR Ab disrupts IFN signaling by preventing released IFN from binding the receptor (30). Thus, one way that σ3 may decrease the sensitivity to this antibody could be through increased synthesis or secretion of Type I IFN. An increased level of type I IFN would be expected to drive greater activation of the JAK-STAT pathway and consequently decrease the sensitivity of the cells to a JAK inhibitor. To determine if levels of IFN-β (the predominant type I IFN expressed by fibroblasts) are altered following σ3 knockdown, we quantified IFN-β mRNA in T3D-infected L929 cells by reverse transcription (RT)-quantitative PCR (qPCR) (Figure 6A). At 18 h following infection, the expression of IFN-β was ∼6-7 fold greater following σ3 knockdown relative to control siRNA-treated cells. To confirm that this increase in IFN-β transcript levels also results in an increase in released protein from cells, we quantified the amount of IFN-β in the media of infected cells by ELISA (Figure 6B). At 24 h following infection, we detected ∼3 fold more IFN-β in the media of infected cells following σ3 knockdown relative to control siRNA-treated cells. Consistent with a greater level of IFN-β, RT-qPCR analysis indicated that in comparison to control siRNA-treated cells, σ3 siRNA-treated cells contained ∼2 fold more mRNA of a representative ISG, ZBP1, at 18 h post infection (Figure 6C). Thus, IFN signaling is more active in cells in which σ3 is knocked down. We have proposed that one or more yet unidentified ISGs are required for the induction of cell death (9). Thus, a decrease in the capacity of the anti-IFNAR antibody to cell death may be related to a diminishment in its efficacy to block ISG expression. To test this idea, we quantified levels of ZBP1 mRNA in T3D-infected cells following treatment with the anti-IFNAR Ab relative to untreated cells using RT-qPCR (Figure 6D). We observe that following treatment with the anti-IFNAR Ab in control-siRNA treated cells, there is a ∼400 fold reduction in the levels of ZBP1 mRNA at 18 h following infection. However, following σ3 knockdown, there is only an ∼80 fold reduction in the levels of ZBP1 mRNA. Thus, σ3 plays a role limiting the induction of IFN-β production following reovirus infection. Further, these data suggest that an increase in IFN-β expression contributes to a greater induction of cell death in reovirus infected cells.

**Figure 6.**
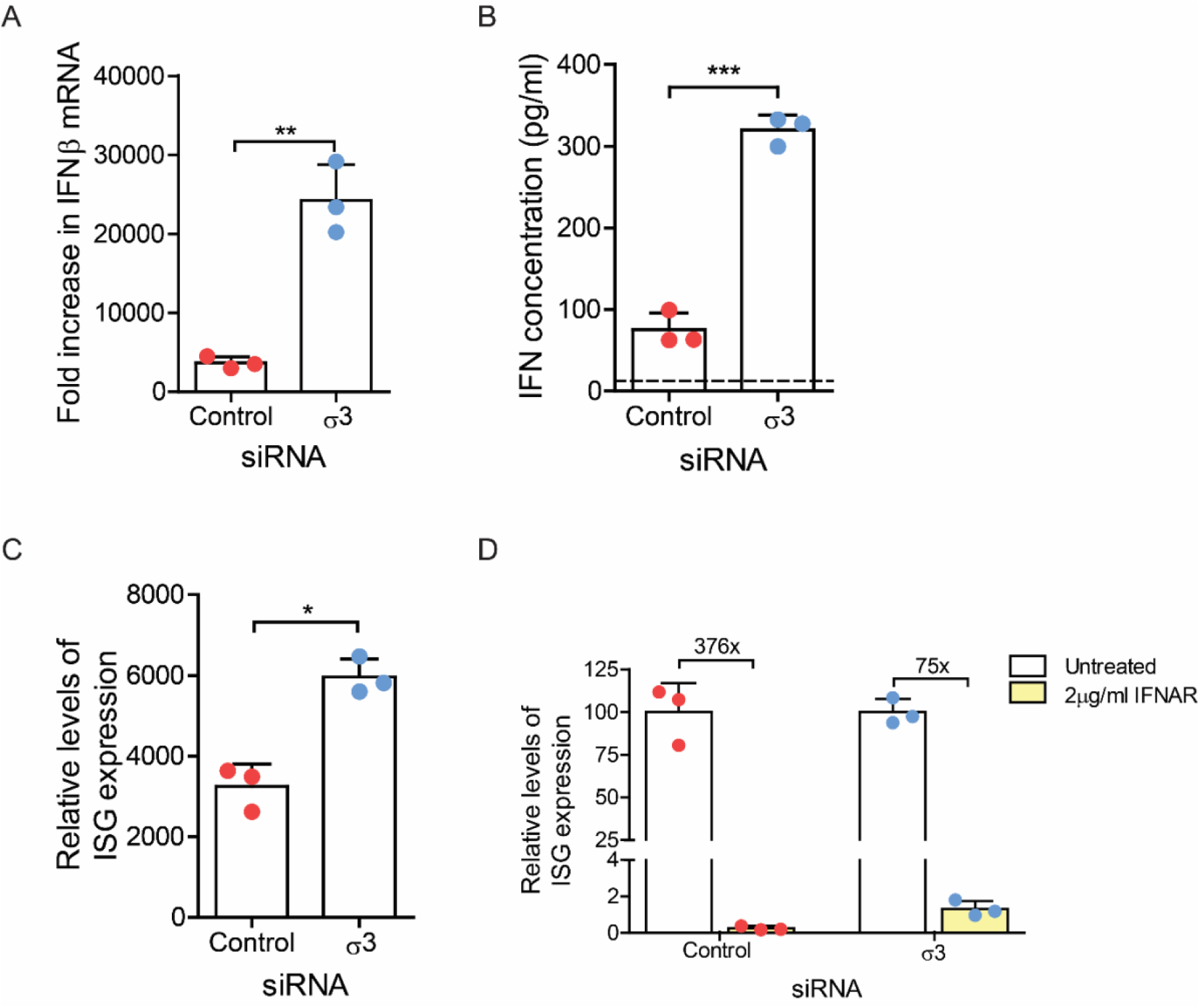
Knockdown of σ3 in L929 cells results in enhanced production of IFN-β and ISGs. L929 cells were transfected with β-gal or σ3 siRNA using INTERFERin. 24 h following transfection, the cells were either mock infected or infected with T3D at an MOI of 10 PFU/cell. (A) RNA extracted from mock or T3D infected cells harvested at 18 h post infection was reverse transcribed using random primers. Fold increase in levels of IFNβ mRNA relative to GAPDH mRNA for T3D infection compared to mock infection was quantified by qPCR and comparative C_T_ analyses. (B) Media harvested from mock or T3D infected cells at 24 h post infection was subjected to an ELISA specific for IFN-β. Data are represented as average concentration of IFN-β in technical duplicates for three independent infections. Dotted line indicates limit of detection (C and D) 24 h following transfection, the cells were either mock infected or infected with T3D at an MOI of 10 PFU/cell. (D) Infected cells were either left untreated or treated with 2µg/ml anti-IFNAR mAb when infection was initiated. (C and D) RNA extracted from mock or T3D infected cells harvested at 18 h post infection was reverse transcribed using random primers. Fold increase in levels of ZBP1 mRNA relative to GAPDH mRNA compared to mock infection was quantified by qPCR and comparative C_T_ analyses. (D) Levels of ZBP1 expression in the absence of treatment was set to 100 for each siRNA-treated sample. Mean values for three independent infections are shown. P values were determined by student’s t-test. *, P<0.05; **, P<0.005; ***, P<0.0005

### The role of σ3 in limiting the type I IFN response is not through its dsRNA binding domain

Induction of IFN by reovirus occurs via detection of incoming dsRNA genome by cellular sensors RIG-I and MDA5 (9). Since these sensors reside in the cytoplasm, this genomic RNA must be exposed to the cytoplasm. The σ3 protein of reovirus contains a dsRNA binding domain (32-34). The dsRNA binding domain of σ3 could allow it to also interact with viral genomic dsRNA in the cytoplasm. Thus, one mechanism by which σ3 could limit induction of IFN-β production is through binding genomic dsRNA and decreasing detection of genome by cellular sensors. To evaluate this possibility, we overexpressed the WT σ3 protein from T3D in HEK293 cells (Figure 7A). The transfected cells were infected with UV-inactivated T3D for 20 h. UV-inactivated virus was used to ensure that no *de novo* gene expression of WT σ3 would occur. We then quantified levels of IFN-β mRNA by RT-qPCR (Figure 7B). Consistent with previous work, UV-inactivated T3D potently induced IFN-β expression (35). Relative to empty vector control, expression of WT σ3 from T3D significantly diminished levels of IFN-β. To evaluate if this effect is produced as a consequence of the capacity of σ3 to bind dsRNA, we determined if a dsRNA binding mutant of σ3 also diminishes UV-inactivated virus induced IFN-β. The K293 residue of σ3 is a previously identified site required for dsRNA binding and the K293T mutation abolishes dsRNA binding (34). We found that σ3 K293T also efficiently blocked UV-virus induced IFN-β expression (Figure 7A, 7B). These data suggest that the mechanism by which σ3 limits IFN induction is not through its dsRNA binding capacity.

**Figure 7.**
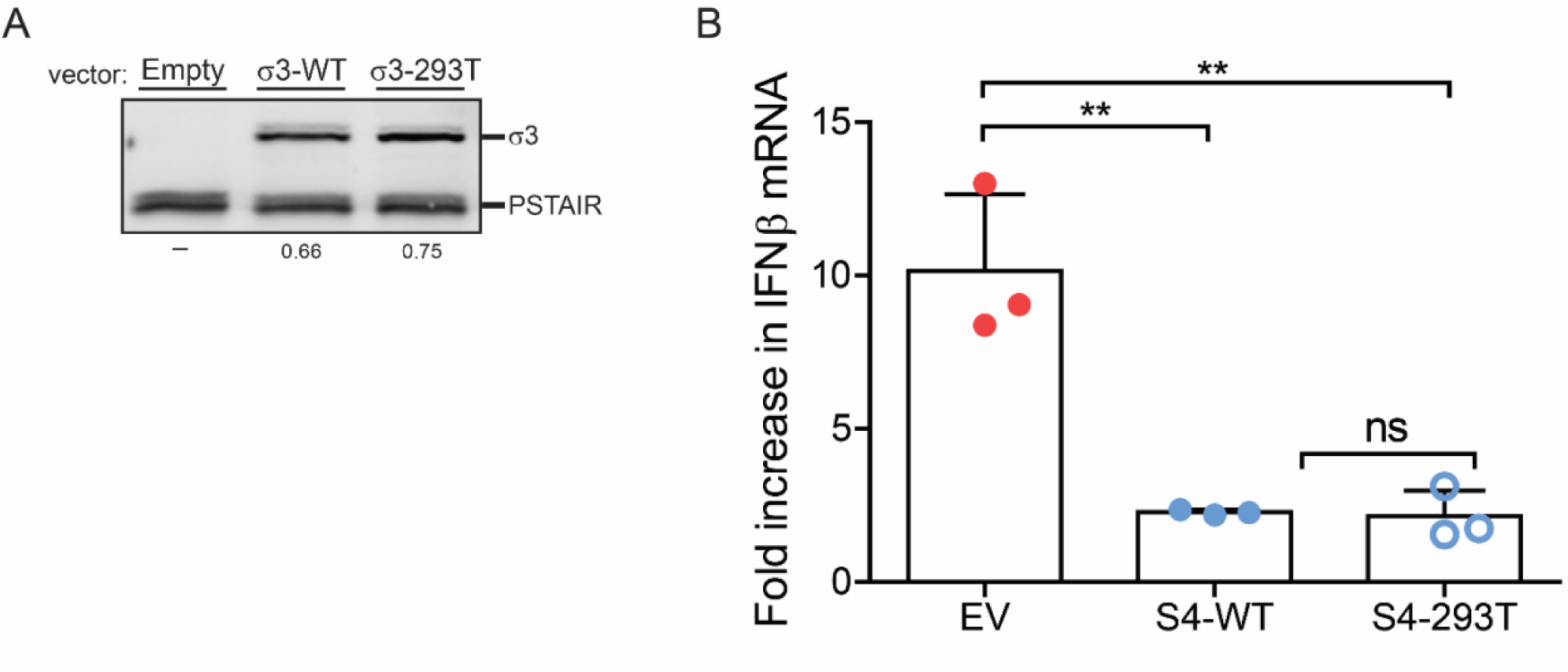
Expression of σ3 in HEK293 cells blocks production of IFN-β following reovirus infection. HEK293 cells were transfected with an empty vector, a vector encoding T3D σ3, or a vector encoding K293T mutant of σ3 using lipofectamine 2000. (A) Cell lysates prepared 48 h following transfection were immunoblotted for the σ3 protein using anti-σ3 4F2 mAb and anti-PSTAIR mAb. Levels of σ3 bands relative to PSTAIR are indicated. (B) 24 h following transfection, cells were either mock infected or infected using UV-inactivated T3D at an MOI of 10 PFU/cell. RNA extracted from mock or UV-T3D infected cells harvested at 18 h post infection was reverse transcribed using random primers. Fold increase in levels of IFNβ mRNA relative to GAPDH for UV-T3D infected cells compared to mock infected cells was quantified by qPCR and comparative C_T_ analyses. Mean values for three independent infections are shown. P values were determined by student’s t-test. *, P<0.05; **, P<0.005; ***, P<0.0005

We demonstrated above that σ3 protein from prototype strain T3D limits IFN-β production and resultant necroptotic cell death. Because prototype reovirus strains display differences in many properties (5), we next examined whether σ3 from prototype strain T1L also can suppress IFN-β production and cell death. In comparison with T3D, T1L induces a lower level expression of IFN-β (36). T1L also induces necroptosis in L929 cells but with slower kinetics (7, 10). Due to sequence similarity of the T3D and T1L S4 mRNA, the siRNA that we have used above also effectively diminishes expression of T1L σ3 (not shown). Analogous to our observations with T3D, we found that knockdown of T1L σ3 enhanced both cell death and IFN-β expression in L929 cells (Figure 8A and 8B). Thus, the function of σ3 appears conserved in both type I and type 3 reovirus strains. We next compared the capacity of WT T1L and a T1L σ3 dsRNA binding mutant to induce the expression of IFN-β and elicit cell death. The R296 residue of σ3 is a previously identified site required for dsRNA binding (34). This positively charged residue, when mutated, no longer maintains affinity for dsRNA. Thus, we utilized a T1L virus that expressing σ3 harboring R296T mutation within the dsRNA binding motif. We found that T1L and T1L S4 R296T induced an equivalent level of IFN-β at 24 h post infection (Figure 9B). Correspondingly, T1L and T1L S4 R296T induced cell death to a similar extent at 48 h post infection with T1L S4 R296T inducing a slightly lower amount than WT T1L (Figure 9C). These data indicate that even in the context of infection, the dsRNA binding capacity of T1L σ3 does not restrict the capacity of the virus to induce IFN-β expression and evoke cell death. Thus, the capacity of σ3 to suppress IFN-β is distinct from its known functions.

**Figure 8.**
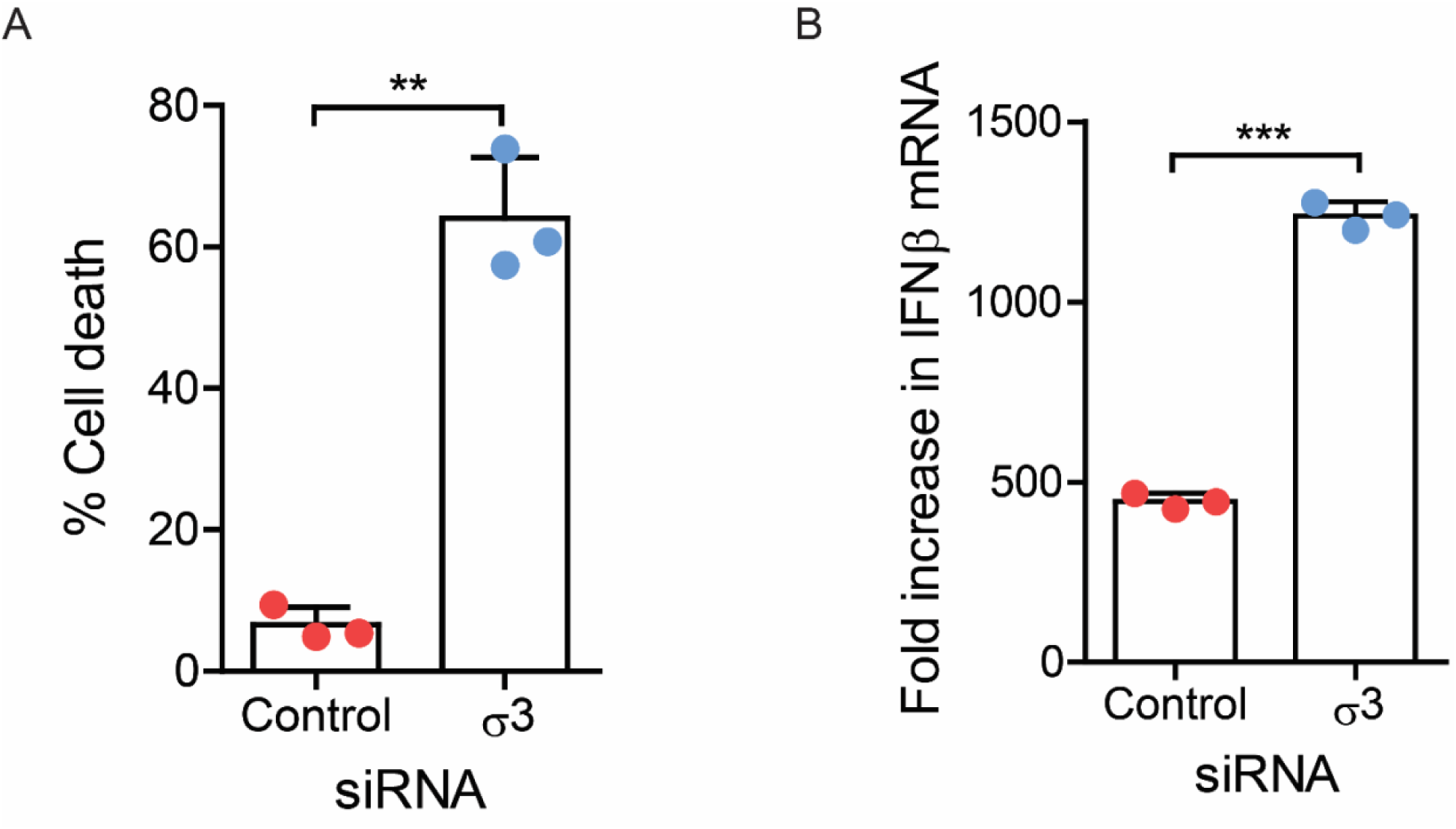
Knockdown of σ3 enhances cell death and IFN-β production following T1L infection. L929 cells were transfected with either β-gal or σ3 siRNA using (A) Lipofectamine 2000 or (B) INTERFERin. (A) 24 h following transfection, the cells were infected with T1L at an MOI of 100 PFU/cell. 24 h following infection, cell death was quantified by AOEB staining. (B) 24 h following transfection cells were either mock infected or infected with T1L at an MOI of 10 PFU/cell. RNA extracted from mock or T1L infected cells harvested at 19 h post infection was reverse transcribed using random primers. Fold increase in the levels of IFN-β mRNA relative to GAPDH for T1L infected cells compared to mock infected cells was quantified by qPCR and comparative C_T_ analyses. Mean values for three independent infections are shown. P values were determined by student’s t-test. **, P<0.005; ***, P<0.0005

**Figure 9.**
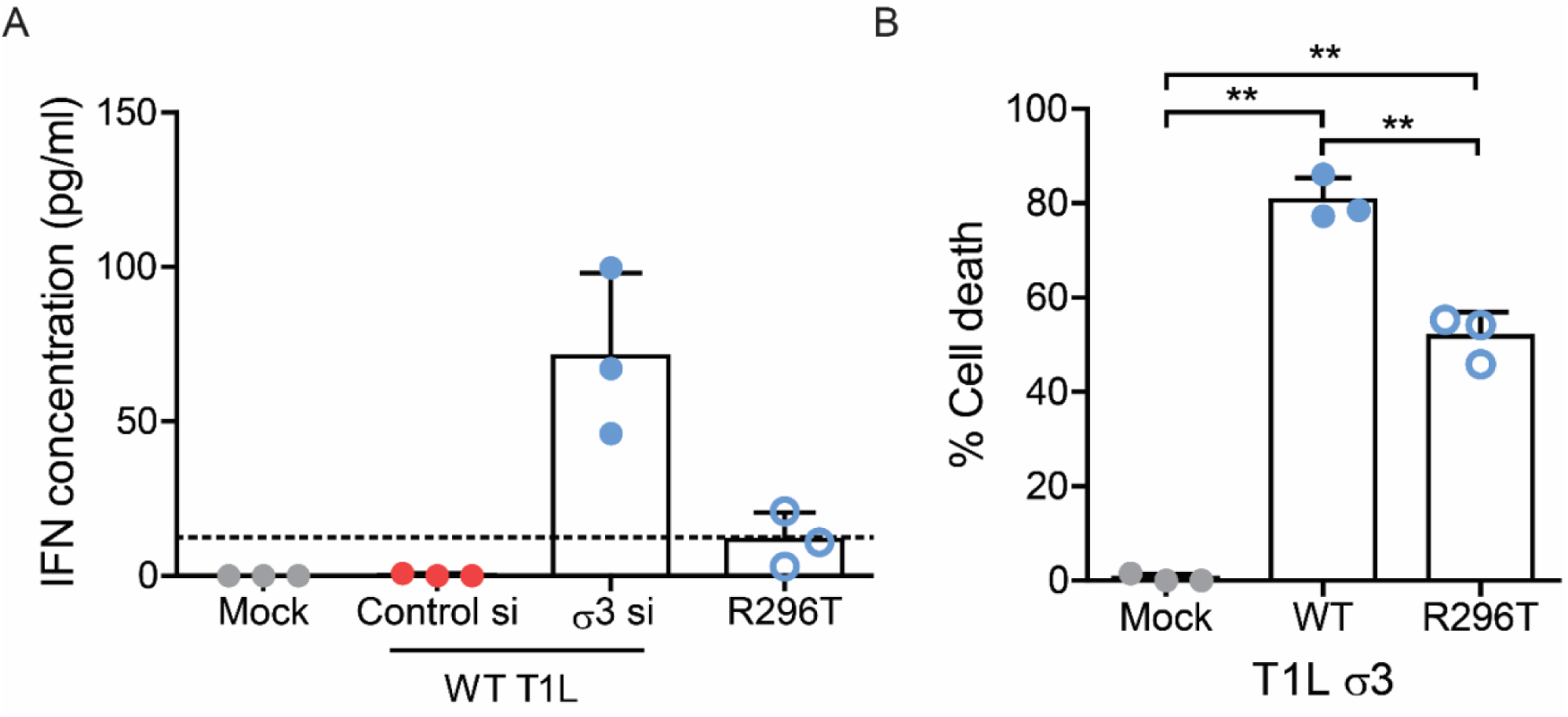
Figure 9. The dsRNA binding function of σ3 does not impact cell death. (A) L929 cells were either left untransfected or transfected with either β-gal or σ3 siRNA using INTERFERin. 24 h following transfection, the cells were either mock infected or infected with WT T1L or T1L-S4-R296T at an MOI of 10 PFU/cell. Media harvested from mock or virus infected cells at 24 h post infection was subjected to an ELISA specific for IFN-β. Data are represented as average concentration of IFN-β in technical duplicates for three independent infections. Dotted line indicates limit of detection (B) L929 cells were either mock infected or infected with WT T1L or T1L-S4_R296T at an MOI of 10 PFU/cell. At 48 h following infection, cell death was quantified by AOEB staining. Mean values for three independent infections are shown. P values were determined by student’s t-test. *, P<0.05; **, P<0.005

## DISCUSSION

We previously demonstrated that knockdown of outer capsid protein µ1 increases necroptosis as a consequence of increased viral gene expression and production of new dsRNA genomes (10). Because σ3 interacts with µ1 to together form the outer capsid, we hypothesized that σ3 may influence necroptosis in a fashion similar to µ1. We report that diminishment in the levels of newly synthesized σ3 does not influence the extent of apoptosis but increases RIP3 dependent necroptosis. However, the mechanism by which σ3 knockdown increases cell death is distinct from that observed following µ1 knockdown. σ3 does not impact levels of viral plus or minus strand RNAs. Instead, σ3 increases the production of IFN-β and consequent downstream IFN signaling (Figure 10). The capacity of σ3 to interfere with IFN-β production appears unrelated to its dsRNA binding property. Thus, our study reveals a new role for σ3 in controlling the host innate immune response.

**Figure 10.**
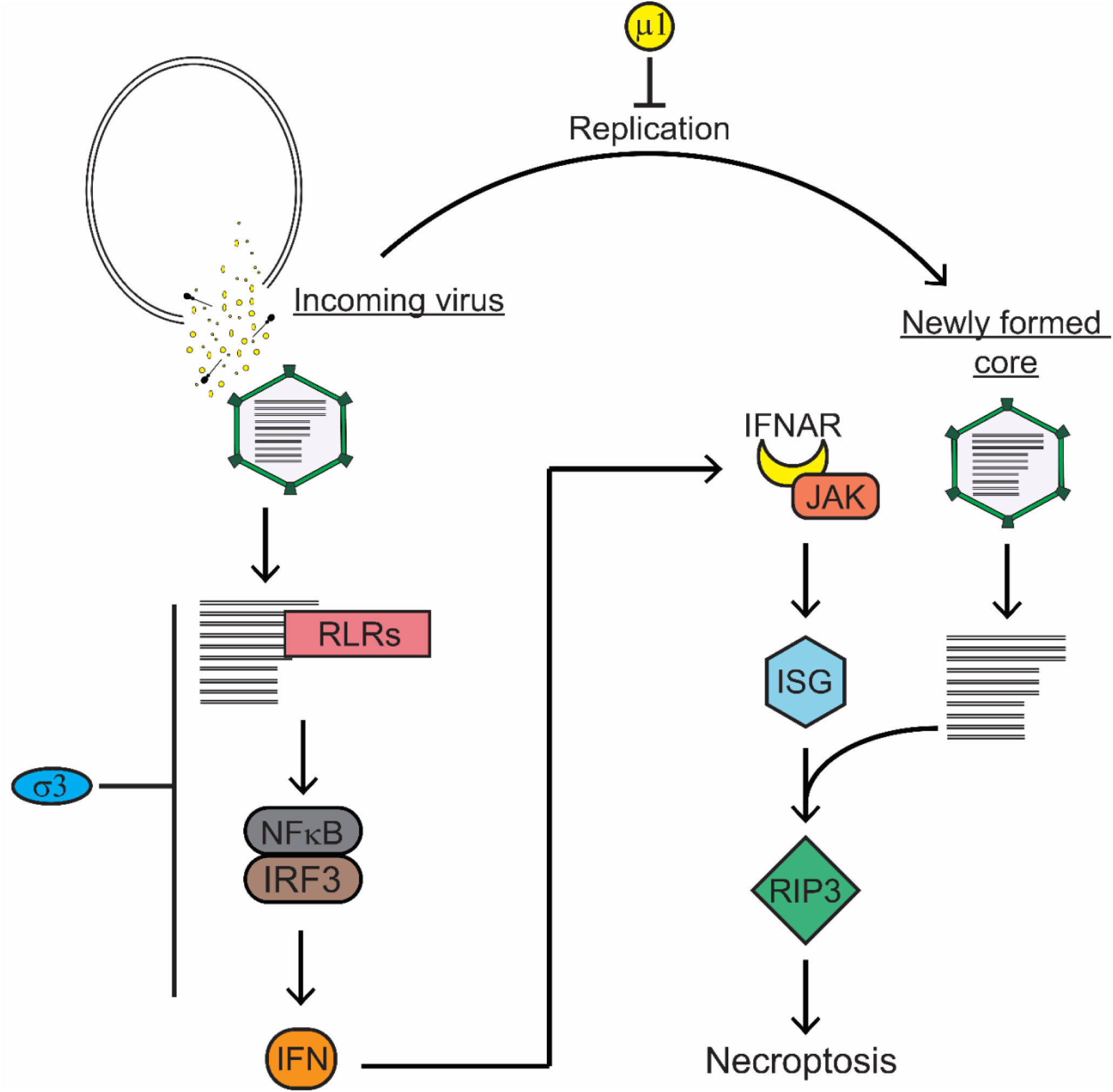
Model depicting the roles of µ1 and σ3 in limiting necroptosis. Pathway of reovirus-induced necroptosis illustrating that in addition to the previously identified role of µ1 in limiting viral replication, σ3 plays an inhibitory role on IFN production and IFN signaling. Therefore, both viral outer capsid proteins negatively regulate the induction of cell death.

Other viral proteins have been implicated in controlling reovirus-induced IFN signaling. µNS is suspected to limit activity of IRF3 by contributing to the formation of viral inclusions which appear to sequester IRF3 and prevent its translocation to the nucleus (37). However, this observation may be cell type specific since our own work using L929 cells (the primary cell type used in this study) indicate that IRF-dependent gene expression is intact in reovirus infected cells (38). Reovirus strains T1L and T3D differ in IFN signaling (36). For example, T1L produces significantly less IFN-β than T3D. The reovirus µ2 protein has been implicated in regulating this difference at least in some cell types (36). This difference has been mapped to a single amino acid polymorphism in µ2 (39). Though the same µ2 polymorphism regulates the morphology of reovirus replication factories (40), how µ2 impacts IFN induction is not clear. The µ2 protein is also suggested to contribute to strain specific differences in sensitivity of reovirus to IFN (36). µ2 properties influence the sequestration of IRF9 in the nucleus (41). Since IRF9 is required for formation of the ISGF3 complex comprised of STAT and IRF9, its sequestration leads to a diminishment in production of ISGs. Our new data demonstrate σ3 plays an inhibitory role on the production of IFN in at least two cell types (Figure 6 and 7). We also demonstrate that σ3 limits IFN production by both reovirus T3D and T1L (Figure 6 and 8). Whether the differences in the capacity of these two reovirus strains to induce IFN-β is related to differences in the properties of σ3 remains to be determined. Future studies using reovirus reassortants or mutants could address this possibility.

It is important to note that viral genome exists in infected cells in two populations – dsRNA genome from incoming viral capsids, and newly synthesized dsRNA genome, made subsequently to viral gene expression (5). Our lab and others have demonstrated that reovirus stimulates the production of IFN through detection of incoming genomic dsRNA (9, 35). UV-inactivated virus, which carries genome but is non-replicative, does not generate new dsRNA or elicit cell death, but still activates IFN signaling (35). Similarly, treatment with ribavirin, which inhibits gene expression, prevents synthesis of new dsRNA but does not impact production of IFN (9). Interestingly, this detection occurs in the cytoplasm via RIG-I and MDA5. Thus, incoming dsRNA must be accessible to these cytoplasmic sensors. Since newly synthesized σ3 limits IFN production by reovirus, we expect that σ3 expression must be concomitant with detection of dsRNA and the attendant host response. One way to explain our observation is that core particles that are delivered into the cytoplasm are initially stable and transcribing viral genes. Over time, these cores fall apart or are degraded, exposing the viral genome and facilitating its detection. If σ3 is expressed prior to this event, it can suppress IFN signaling.

Another possible explanation for our observations would relate to the high particle-to-PFU ratio (∼ 100) of reovirus (5). It is possible that some incoming viral particles are degraded during the entry process and expose the viral genome whereas the intact particles express viral gene products such as σ3 and suppress IFN signaling. This scenario would also require that σ3 expression precede degradation of non-infectious particles. Our ongoing work is focused on elucidating how the genome of the incoming virus particles is exposed.

It is unclear how the roles of σ3 and µ1 in necroptosis coordinate temporally. Other roles for these proteins have been described including µ1’s role in activating apoptotic signaling and σ3’s role in promoting viral translation (19,22, 23, 42, 43). These two outer capsid proteins also interact late in infection to form heterohexamers on the newly made viral particles (13). This interaction disrupts their individual functions (23, 42). Since σ3 and µ1 coassemble on progeny particles, it is unlikely that σ3-µ1 interaction blocks the function of µ1 in controlling the accumulation of viral gene products. However, no data exist to directly support this idea. Whether µ1 impacts the IFN suppression function of σ3 also remains unknown. The affinity between µ1 and σ3 differs between strains of reovirus and this difference in affinity can influence the degree to which µ1 could affect σ3 function to limit the IFN response (23). Furthermore, this could account for strain specific differences in IFN induction.

σ3 is known to be an antagonist of PKR (19, 23). PKR is activated by binding to dsRNA (21). Once activated PKR phosphorylates eIF2α leading to a shut off of host and viral translation. By binding dsRNA, σ3 prevents PKR activation, thereby increasing viral translation. Consistent with this idea, we observed that decreasing σ3 levels results in diminished expression of other viral proteins (not shown). These other proteins may have their own effect on IFN signaling. Therefore, the increased production of IFN-β that we observe following σ3 knockdown may be due to an indirect effect of σ3’s role in promoting the translation of other viral proteins. However, since overexpression of σ3 in HEK293 cells is alone sufficient to block IFN-β expression, we think it is more likely that this effect is direct. Nonetheless because dsRNA binding mutants of σ3 that cannot prevent PKR activation also can block IFN-β expression, these data suggest that blockade of protein synthesis by σ3 knockdown occurs via a mechanism that is independent from its effect on PKR function. We propose that diminished protein synthesis following σ3 knockdown may be a downstream consequence of expression another ISG that blocks viral translation.

The capacity for reovirus to induce cell death is the basis for many aspects of its biology including both its pathogenesis and its oncolytic potential. Our data demonstrate that σ3 plays a role in limiting the innate immune response and cell death. Therefore, this study provides a preliminary basis by which we can manipulate the cellular response to infection.

